# A multidrug resistant clinical P. aeruginosa isolate in the MLST550 clonal complex: uncoupled quorum sensing modulates the interplay of virulence and resistance

**DOI:** 10.1101/415000

**Authors:** Huiluo Cao, Tingying Xia, Yanran Li, Zeling Xu, Salim Bougouffa, Yat Kei Lo, Vladimir B. Bajic, Haiwei Luo, Patrick C.Y. Woo, Aixin Yan

**Affiliations:** School of Biological Sciences, The University of Hong Kong, Pokfulam Road, Hong Kong SAR, China; Computational Bioscience Research Center (CBRC), King Abdullah University of Science and Technology (KAUST), Thuwal, Saudi Arabia; School of Life Sciences, The Chinese University of Hong Kong, Sha Tin, Hong Kong SAR, China; Department of Microbiology, Li Ka Shing Faculty of Medicine, The University of Hong Kong, Hong Kong SAR, China

## Abstract

*Pseudomonas aeruginosa* is a prevalent and pernicious pathogen equipped with both extraordinary capabilities to infect the host and to develop antimicrobials resistance (AMR). Monitoring the emergence of AMR high risk clones and understanding the interplay of their pathogenicity and antibiotic resistance is of paramount importance to avoid resistance dissemination and to control *P. aeruginosa* infections. In this study, we report the identification of a multidrug resistant (MDR) *P. aeruginosa* strain PA154197 isolated from a blood stream infection in Hong Kong. PA154197 belongs to a distinctive MLST550 clonal complex shared by two international *P. aeruginosa* isolates VW0289 and AUS544. Comparative genome and transcriptome analysis with the reference strain PAO1 led to the identification of a variety of genetic variations in antibiotic resistance genes and the hyper-expression of three multidrug efflux pumps MexAB-OprM, MexEF-OprN, and MexGHI-OpmD in PA154197. Unlike many resistant isolates displaying an attenuated virulence, PA154197 produces a significantly high level of the *P. aeruginosa* major virulence factor pyocyanin (PYO) and displays an uncompromised virulence compared to PAO1. Further analysis revealed that the secondary quorum sensing system Pqs which primarily controls the PYO production is hyper-active in PA154197 independent of the master QS systems Las and Rhl. Together, these investigations disclose a unique, uncoupled QS mediated pathoadaptation mechanism in clinical *P. aeruginosa* which may account for the high pathogenic potentials and antibiotics resistance in the MDR isolate PA154197.

## Introduction

*Pseudomonas aeruginosa* is a ubiquitous Gram-negative pathogen that causes a variety of notorious infections in humans such as ventilator-associated pneumonia, lung infections of cystic fibrosis (CF) patients, burn wound infection, and various sepsis syndromes. It is the second leading cause of hospital-acquired infections and is especially problematic in ICUs, where it is the leading cause of pneumonia among pediatric patients and is responsible for a large number of urinary tract (10% in US and 19% in Europe), blood stream (3% in US and 10% in Europe), eye, ear, nose, and throat infections (1-3). Compounding the burden of these infections is the extraordinary capability of the pathogen to develop antibiotic and multidrug resistance (MDR) even during the course of antibiotics therapy. *P. aeruginosa* is one of the “ESKAPE” (*Enterococcus spp. Staphylococcus aureus Klebsiella spp. Acinetobacter baumannii Pseudomonas aeruginosa*, and *Enterobacter spp*.) organisms which are recognized by the WHO as an alarming threat to the global public heath associated with antimicrobial resistance (AMR). As a consequence, the diseases outcome of the *P. aeruginosa* infections is the complex interplay of the pathogen (its pathogenicity and virulence), hospital environments (antibiotic therapies and the emergence of AMR), and the patient’s conditions (host immune responses) (4-6).

*P. aeruginosa* is genetically equipped with outstanding intrinsic antibiotic resistance machineries. These include the inducible production of the AmpC cephalosporinase, presence of the housekeeping MexAB-OprM multidrug efflux pump, and the limited permeability of its outer membrane caused by low expression and inefficient porin proteins (4, 7, 8). In addition, the pathogen has extraordinary capabilities of developing acquired antibiotic resistance. Mutational overexpression of one of at least four efflux pumps, MexAB-OprM, MexCD-OprJ, MexEF-OprN, and MexXY-OprM, encoded in the genome of *P. aeruginosa* often plays an important role in acquired resistance and can lead to clinically significant MDR (9, 10). Among them, the MexAB-OprM pump displays the broadest substrates profile and mutational overexpression of this pump can lead to resistance to all β-lactams (except imipenem), (fluoro)quinolones, tetracyclines, and macrolides in clinics. Similar to the MexAB-OprM pump, overexpression of MexXY is frequent (10-30%) among clinical strains and causes decreased susceptibility to aminoglycosides and cefepime. Overexpression of MexEF-OprN and MexCD-OprJ is less frequent in clinical isolates (<5%) and they mainly affect fluoroquinolones resistance (5, 11). In addition to efflux pump overexpression mediated MDR, *P. aeruginosa* also readily develops resistance to specific class of antibiotics through genetic mutations and acquisition of target or drug inactivation genes. These include overexpression of *ampC* caused by mutations in peptidoglycan-recycling genes *ampD, dacB*, or *ampR* that causes resistance to noncarbapenem β-lactams; acquisition of aminoglycosides modification enzymes (AMEs) and ribosomal methyltransferase (Rmts) enzymes which are associated with aminoglycosides resistance; mutations in the DNA gyrase (*gyrA* and *gyrB*) and/or topoisomerase IV (*parC* and *parE*) which results in fluoroquinolone resistance, and mutational repression of the OprD porin which leads to resistance to imipenem (12-15).

Regardless of the resistance mechanisms involved, the emergence and prevalence of MDR *P. aeruginosa* strains continue to rise rapidly world-wide. Although the bacterium displays an overall nonclonal epidemic population structure with most isolates represented by single MLST (multilocus sequence typing) genotypes, MDR or XDR (extensively drug-resistant) *P. aeruginosa* isolates display a much lower clonal diversity than the susceptible isolates and recent studies have reported the existence of MDR/XDR global clones disseminated in different hospitals world-wide (11, 16, 17). Inter-patient spread of antibiotic resistant mutations linked to the transmission of epidemic CF strains is also reported. These clones are denominated as international high-risk clones. Among them, the most recognized successful clones are the high risk MDR clones ST111, ST175, and ST235, and the Liverpool Epidemic Strain (LES) ST146, which is the most epidemic clone among CF patients (11). Further deeper analysis of the molecular details of their resistance development and closely monitoring of the emerging of new international MDR/XDR high risk clones are of paramount importance to avoid the world-wide dissemination of these clones and to control *P. aeruginosa* infections.

In addition to their broad and high level of antibiotic resistance, many MDR clones, especially the three major international high risk clones ST111, ST175, ST235, are found to be associated with a defined set of biological markers which include defective motility (swimming, swarming, and twitching), reduced pigment production (pyocyanin and pyoverdine), reduced in vitro fitness, but enhanced biofilm formation (5, 11). These suggest the presence of a fitness trade-off that compromises the pathogenic potentials and virulence of MDR isolates. Several epidemiological survey and molecular evolutionary studies ascribed these phenotypes to quorum sensing (QS) deficiency in these strains (18-21), which is not unexpected since production of many virulence factors, such as exo-proteases, elastase, rhamnolipids, lectin, pyoverdine, pyocyanin, hydrogen cyanide etc. are primarily regulated by QS systems in *P. aeruginosa*(1). On the other hand, recent studies also reported antibiotic resistant strains displaying enhanced virulence (17, 22-24), such as *P. aeruginosa* strains lacking the *oprD* porin (22), implying that certain MDR clones may develop compensatory mutations allowing them to recover their virulence without affecting the level of resistance.

Recently, we surveyed 84 *P. aeruginosa* clinical strains isolated from various of infections including wound (both superficial and deep), urine, ear, pus, drain fluid, and blood in the Queen Mary Hospital, Hong Kong, and identified an MDR isolate PA154197 from a blood stream infection. PA154197 displays an MDR level and profile comparable to the international high-risk clone ST175. Sequencing its genome reveals that it belongs to a distinctive genotype MLST550 rather than the ST175 clone. The MLST550 genotype is shared by two additional clinical strains in the database: VW0289 and AUS544 which were isolated from the sputum and bronchial lavage of CF patients in The Netherlands and Australia, respectively, suggesting a potential international transmission of this emerging clonal complex. This combined with the extraordinary antibiotic resistance profile of the strain prompt us to use PA154197 as a prototype to conduct comparative genomic and transcriptomic analysis and understand the resistance development and virulence of the MLST550. These studies led to the identification of the adaptive activation of the secondary quorum sensing system Pqs (*Pseudomonas* quinolone system) independent of the primary QS system Las and Rhl which may account for the uncompromised virulence in the MDR strain PA154197. Together, our studies report the emergence of a high-risk clone in clinics and its underlying pathoadapation mechanism mediated by uncoupled QS.

## Results

### PA154197 displays extraordinary antibiotic resistance

We examined the susceptibility of PA154197 to various classes of antibiotics including the common antipseudomonal drugs aztreonam (ATM), imipenem (IPM), ciprofloxacin (CIP), levofloxacin (LVX), and amikacin (AMK) by measuring their MIC values and compared with that for PAO1 (Fig. 1A). We found that PA154197 displays resistance to almost all these different classes of antibiotics with a 4-, 16-, 80-, and 160-fold decreases in susceptibility to the common antipseudomonal drugs ATM, IPM, CIP, and LVX, respectively, compared to PAO1. The only two antibiotics it remains susceptible are the aminoglycoside amikacin and the cyclic non-ribosomal polypeptides polymyxin B and E, which meets the XDR criteria (resistance to all but one or two of the eight classes of antipseudomonal drugs) according to Magiorakos et. al (16). To further evaluate its resistance level, we compared the antibiotics susceptibility profile of PA154197 with the three international high-risk clones using the published data (25) and found that PA154197 displays a comparable resistance profile with the high-risk clone ST175 (Fig. 1B), highlighting the high epidemic and risk potential of PA154197.

**Figure 1.**
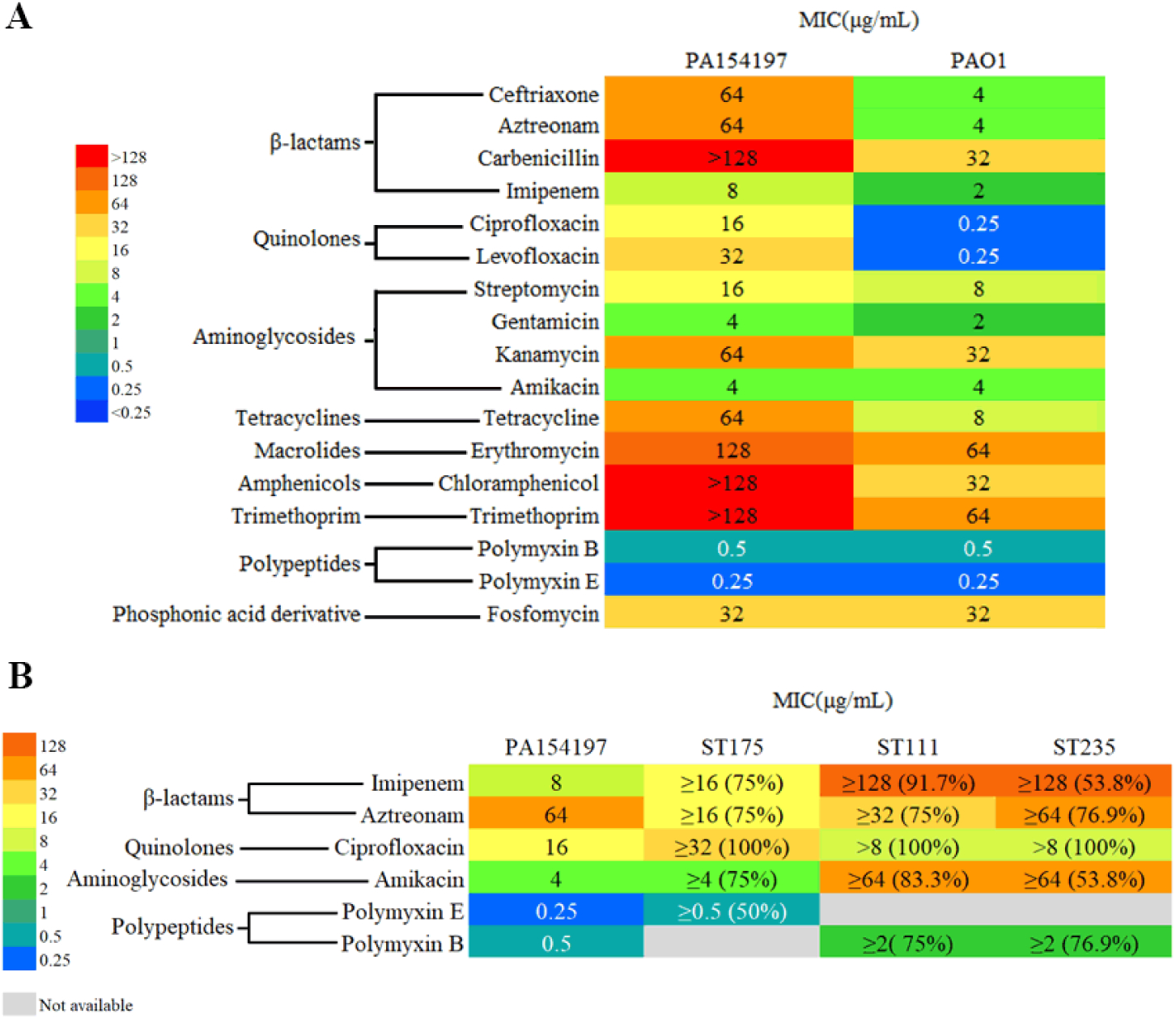
Antibiotic resistance profiles of PA154197. **A.** MIC values of various antibiotics to PA154197 and PAO1. **B.** MIC values of indicated antibiotics to the international high risk clone ST175 are retrieved from published data (25) and are compared with that of PA154197. Resistance levels are displayed in color scheme with low susceptibility (high MIC values) in red color and high susceptibility (low MIC values) in blue color. The color scheme is constructed using background filling application in the Excel of Microsoft Office.

### Sequencing type, phylogenetic position, and genomic islands of PA154197

We then sequenced the genome of PA154197 which resulted in one single circular chromosome with a length of 6,445,239 bp and a GC content of 66.38%. A total number of 5,923 genes are predicted in PA154197 genome which include 5,816 coding gene sequences (CDS), 26 pseudo-genes, 12 rRNA, 65 tRNA and 4 ncRNA (Table 1). Based on PubMLST, PA154197 is assigned to the multilocus sequence typing (MLST) 550, a clonal complex which is currently shared by two additional clinical isolates, VW0289 and AUS544 from the CF patients in the Netherland and Australia, respectively (Fig. 2A), but the complete genome sequences of these two strains are unavailable. PAst analysis revealed that PA154197 belongs to the Serotype O9 (26) (Table 1). Analyzing its accessory genome identified 21 genome islands (GIs) including 3 prophages (Table S1). In addition to hypothetical proteins, a large number of metabolic enzymes such as oxidoreductase, oxygenase and halogenase, as well as the AraC and LysR type transcription regulators are encoded in these GIs, implying the potential roles of the accessory genome of PA154197 in the adaptation and fitness of the bacterium.

**Table 1.** The genomic features of *P. aeruginosa* PA154197.

| Features | Length<br>(bp) | % GC<br>content | Genes | Proteins | rRNA | tRNA | ncRNA | Pseudo<br>genes | CRISPR<br>arrays |
| --- | --- | --- | --- | --- | --- | --- | --- | --- | --- |
| PA154197 | 6,445,239 | 66.4% | 5,923 | 5,816 | 12 | 65 | 4 | 26 | 3 |

**Figure 2.**
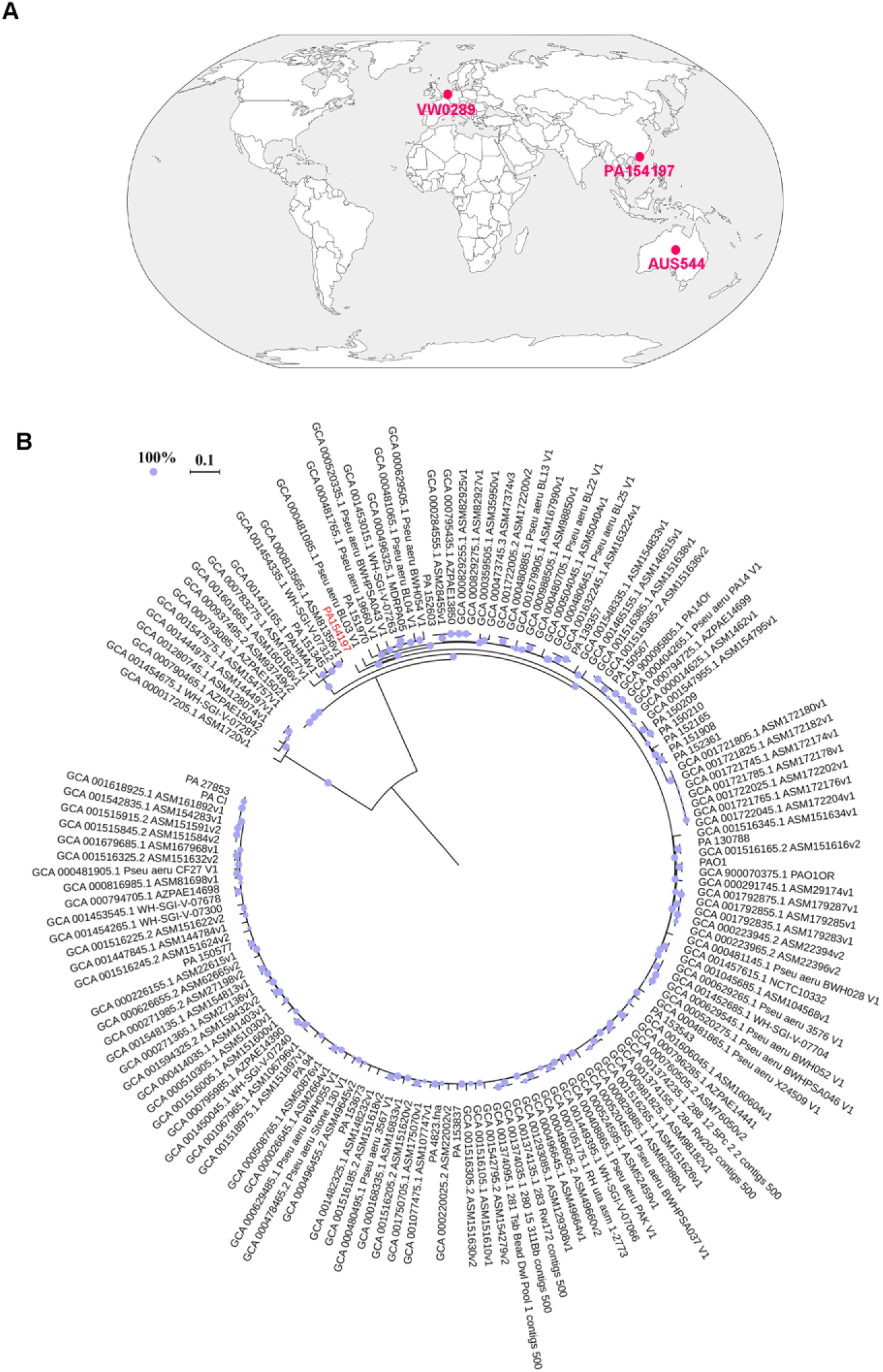
Lineage complex of the MLST550 and phylogenetic position of PA154197. **A.** Geographical distribution of the three MLST550 *P. aeruginosa* strains currently available in pubMLST database, VW0289, VW0289, and PA154197. **B**. phylogenetic relationships of PA154197 with 149 *P. aeruginosa* strains including 78 complete and 71 incomplete genomes from representative 32 groups available in Genbank database (as of October 2017). The phylogenetic tree was constructed based on the SNPs identified using Harvest with 100 bootstrap and maximum likelihood (ML) criterion in MEGA 7 software. Scale bar indicates the branch length and circles on the nodes represent the statistical supports according to size. The denotation of the strain is listed in the Table S3 and PA15419 is highlighted in red.

To examine the phylogenetic relationship of PA154197 with other *P. aeruginosa* strains, we constructed a phylogenetic tree based on 140,634 SNPs generated from 150 *P. aeruginosa* genomes which include all publicly-available complete genomes (78 as of October 2017) and representative non-complete genomes. As shown in Fig 2B, PA154197 exhibits a distinct phylogenetic position from other genomes available in the database. This may indicate its unique virulence and pathogenic potentials.

### Antibiotic resistance genes and mutational changes therein

Using Resistance Gene Identifiers (RGI), we then identified antibiotic resistance genes in PA154197 against the Comprehensive Antibiotic Resistance Database (CARD)(27), and examined genomic variations including mutations, deletions and indels in these genes with the genome of PAO1 as the reference (Table 2). It was revealed that PA154197 contains a large number of mutations associated with antibiotic resistance genes, including both previously reported genetic variations and those newly identified in PA154197. Several well established mutations include an 8-bp deletion in the *mexT* gene which leads to up-regulation of the MexEF-OprN efflux pump and down-regulation of the OprD porin, and consequently resistance to fluoroquinolones and imipenem; the T105A substitution in AmpC, which causes resistance to non-carbapenem β-lactams (28), and the T83I substitution in the so called quinolone resistance-determining region (QRDR) of GyrA, conferring quinolone resistance. Another genomic variation commonly found to cause resistance in many *P. aeruginosa* isolates is the non-synonymous mutations in the TetR transcriptional regulator NalC which often leads to aztreonam resistance (29). Three non-synonymous mutations, G71-E, E153-Q, S209-R, are identified in the *nalC* gene in PA154197 (Table 2).

**Table 2.** Mutational changes of genes conferring antibiotic resistance and their expression levels in PA154197 relative to that in PAO1.

| Genes | Non-synonymous/Indel mutations in<br>encoding region | Log2FoldChange<br>(PA154197/PAO1) |
| --- | --- | --- |
| *mexS* | D249-N | 1.38 |
| *mexT* | Del (8bp:237-244),F172-I,G301-C | 0.60 |
| *mexE* | N/A | 6.93 |
| *mexF* | N/A | 6.61 |
| *oprN* | N/A | 6.69 |
| *mexG* | N/A | 3.65 |
| *mexH* | N/A | 3.80 |
| *mexI* | A782-E, A936-T | 3.87 |
| *opmD* | N/A | 3.77 |
| *armR* | S21-T,Y32-C |  |
| *mexR* | E76-* | 2.54 |
| *mexA* | N/A | 3.47 |
| *mexB* | I1001-T | 3.52 |
| *oprM* | N/A | 3.44 |
| *mexX* | Q4-H,K329-Q,L331V,W358-R | - |
| *mexY* | T543-A,L984-F | - |
| *mexZ* | N/A | - |
| *mexC* | P57-S,R76-Q,H310-R,S330-A,A378-T,P383-S,A384-V | 2.05 |
| *mexD* | T87-S,A155-T,E257-Q,E314-<br>D,V434-A,N669-D,S685-G,I703-<br>V,S845-A,S915-A,I982-V,K1031- | 0.72 |
|  | R,S1040-T |  |
| *oprJ* | T376-S,K413-M | - |
| *mexJ* | F13-L,A314-P | 2.86 |
| *mexK* | I21-L,K694-R,A841-T | 3.16 |
| *mexL* | S4-P | 0.97 |
| *muxA* | T261-A | - |
| *muxB* | N/A | - |
| *muxC* | N/A | - |
| *opmB* | N/A | - |
| *mexP* | N/A | 1.03 |
| *mexQ* | I294-V,V384-I,G505-D,V517-A,R656-K,D722-E | - |
| *opmE* | S46-G,I50-V,S175-T,W354-R | - |
| *oprD* | D43-N,S57-E,S59-R,E202-Q,I210-A,E230-K,S240-T,N262-T,A267-S,A281-G,K296-Q,Q301-E,R310-G,V359-L,Del(6bp:1132-1137),M372-V,S373-D,D374-S,N375-S,N376-S,V377-S,K380-Y,N381-A,Y382-G,G383-L |  |
| *ampC* | T105-A | -0.88 |
| *ampR* | N/A | - |
| *gyrA* | T83-I, Del(6bp:2734-2739) | - |
| *gyrB* | N/A | - |
| *parC* | N/A | - |
| *parE* | N/A | 0.84 |
| *parS* | H398R | 0.64 |
| *parR* | N/A | 0.61 |
| *mvaT* | N/A | -0.48 |
| *poxB* | N/A | -2.55 |
| *nalC* | G71-E,E153-Q,S209-R | 0.42 |
| *nalD* | N/A | - |
| *aph* | A14-S | - |
| *amgS* | I260-V,G288-D | - |
| *spuF* | P276-S | - |
| *fusA1* | I186-V | 0.89 |
| *fusA2* | S176-A,A690-S,G695-A | -2.78 |

Notes: N/A, no non-synonymous mutation occurred; Ins: insertion; Del: deletion; “-” indicates not significant difference. “\*”: stop codon.

New genomic variations that potentially lead to antibiotic resistance is also identified in PA1541197, such as a 6-bp deletion (corresponding to the deletion of E959 and S969) in the C-terminal of GyrA and a E76-*(stop codon) mutation in the *mexR* gene which will lead to pre-mature termination of MexR at E76 and potentially de-repression of the MexAB-OprM major efflux pump, causing multidrug resistance(30). Mutations that potentially de-repress or induce the expression of several other efflux pumps are also identified in the corresponding transcription regulators or the promoter region of the efflux genes. For example, a single point mutation H398R is identified in *parS* gene which regulates the expression of the *mexEF*-*oprN* efflux pump (31). A 5-nucleotides alteration in the promoter region (−1 to −250bp) of the *mexGHI-opmD* operon is present in PA154197 compared with that in PAO1, which may lead to the expression change of the MexGHI-OpmD efflux pump. A complete summary of the identified genomic variations in all the antibiotic resistance genes is provided in Table 2.

### Expression of the resistance genes and the functional genome of P. aeruginosa PA154197

To examine the resistance mechanisms of PA154197 especially the expression levels of the resistance genes identified, we performed a comparative transcriptome analysis on PA154197 and the reference strain PAO1 using RNA-seq. Among the 5,543 orthologous genes identified in the two strains using progressiveMauve (32), differential expression of 3,148 genes are observed and their expression levels are compared using DESeq (33) (Fig. 3 and Table S2). Notably, among the genes which display significantly higher expression levels in PA154197 than in PAO1, those encoding four efflux pumps, such as *mexEF-oprN* (PA2493-PA2495), *mexRAB-oprM* (PA0424-PA0427), *mexGHI-opmD* (PA4205-PA4208), and *mexKJL* (PA3676-PA3678) are observed (Fig. 3), consistent with the MDR profile of PA154197. Genes that confer resistance to specific antibiotics, such as that of *ampC, gyrA, gyrB*, and *parC parE*, are not differentially expressed in the RNA-Seq analysis, suggesting that over-expression of several multidrug efflux pumps may play a major role in the MDR development in PA154197. COG functional distribution analysis of the differentially expressed genes also reveals the enrichment of efflux pump genes among all the genes which are expressed at a higher level in PA154197 than in PAO1 (Fig S1).

**Figure 3.**
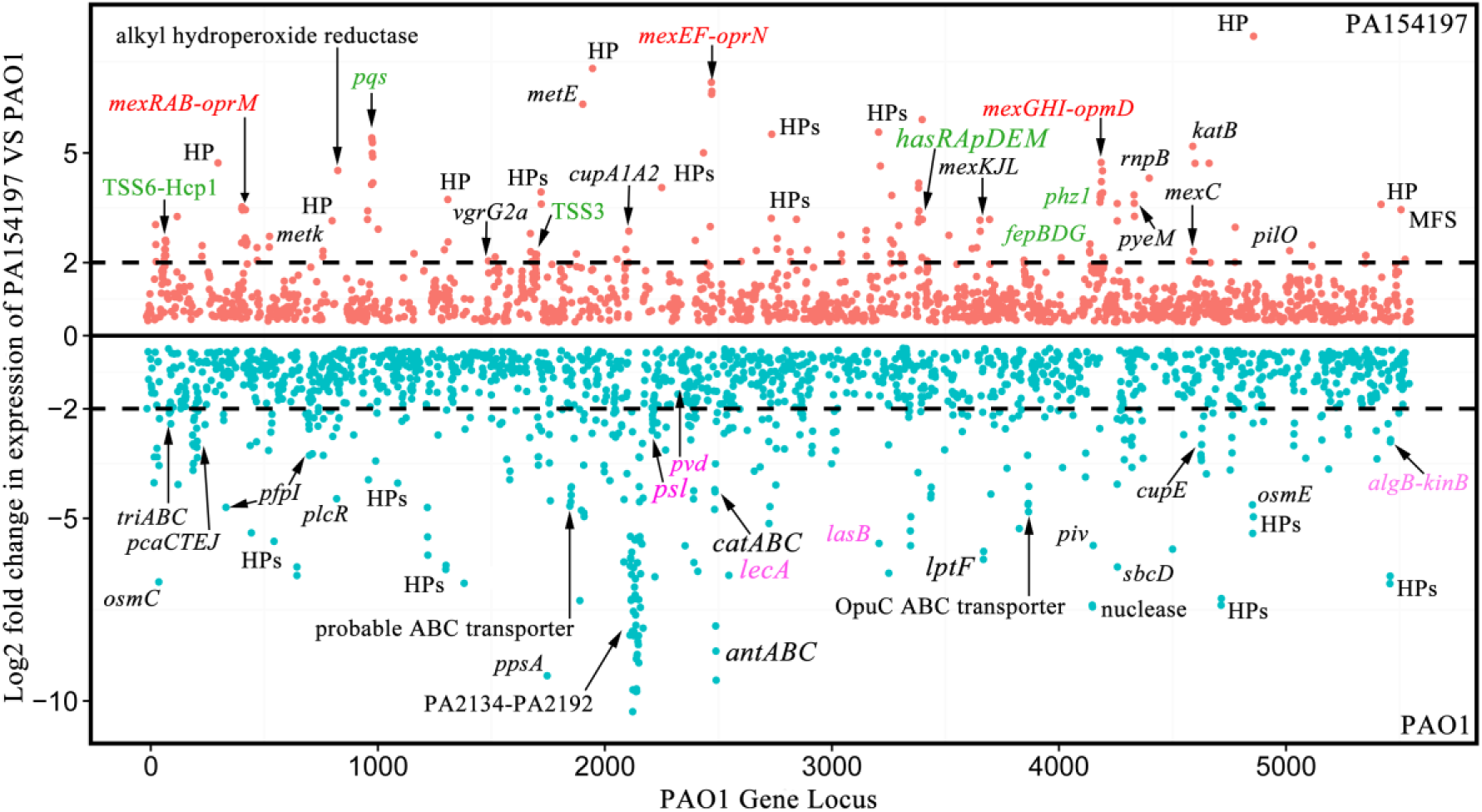
The genome wide transcriptomic profile of *P. aeruginosa* PA154197 and PAO1. Orange dots indicate genes with higher relative expression levels in PA154197 than in PAO1 and blue dots represent genes with higher relative expression level in PAO1 than in PA154197. The black dashed lines represent 4 fold (value of log2 > 2) changes in expression. Genes and operons with distinctive expression patterns in the two strains are indicated. Among them, antibiotic resistance genes are highlighted in red, genes encoding virulence factors are highlighted in green (expressed in a higher level in PA154197 than PAO1) or purple (expressed in a higher level in PAO1 than in PA154197). X-axis represents the gene locus with PAO1 genome as reference.

To verify the relative expression levels of these genes in the two strains, we conducted RT-qPCR analysis. As shown in Fig 4B, consistent with the RNA-Seq data, the MexEF-OprN efflux system displayed the most significant higher expression (265-574 fold) in PA154197 than in PAO1, followed by the MexGHI-OpmD (8.5-41.9 fold) and the MexAB-OprM (4.6-11 fold) efflux system. On the other hand, another efflux system which over-expression is associated with aminoglycosides resistance, the MexXY-OprD system, did not show increased relative expression in PA154197 in either RNA-Seq or the RT-qPCR analysis, consistent with the fact that PA154197 remains susceptible to the aminoglycoside antipseudomonal drug amikacin (Fig 1). Overall, an EB efflux assay showed the hyper-efflux activity in PA154197 than in PAO1 (Fig 4C), suggesting that over-expression of the efflux pumps MexEF-OprN, MexAB-OprM, and MexGHI-OpmD may play a major role in conferring MDR in PA154197. RT-qPCR analysis also revealed the decreased expression of another resistance gene *oprD* in PA154197 which serves as the entry portal of imipenem, consistent with the observed imipenem resistance (MIC as 8μg/ml) in this strain.

**Figure 4.**
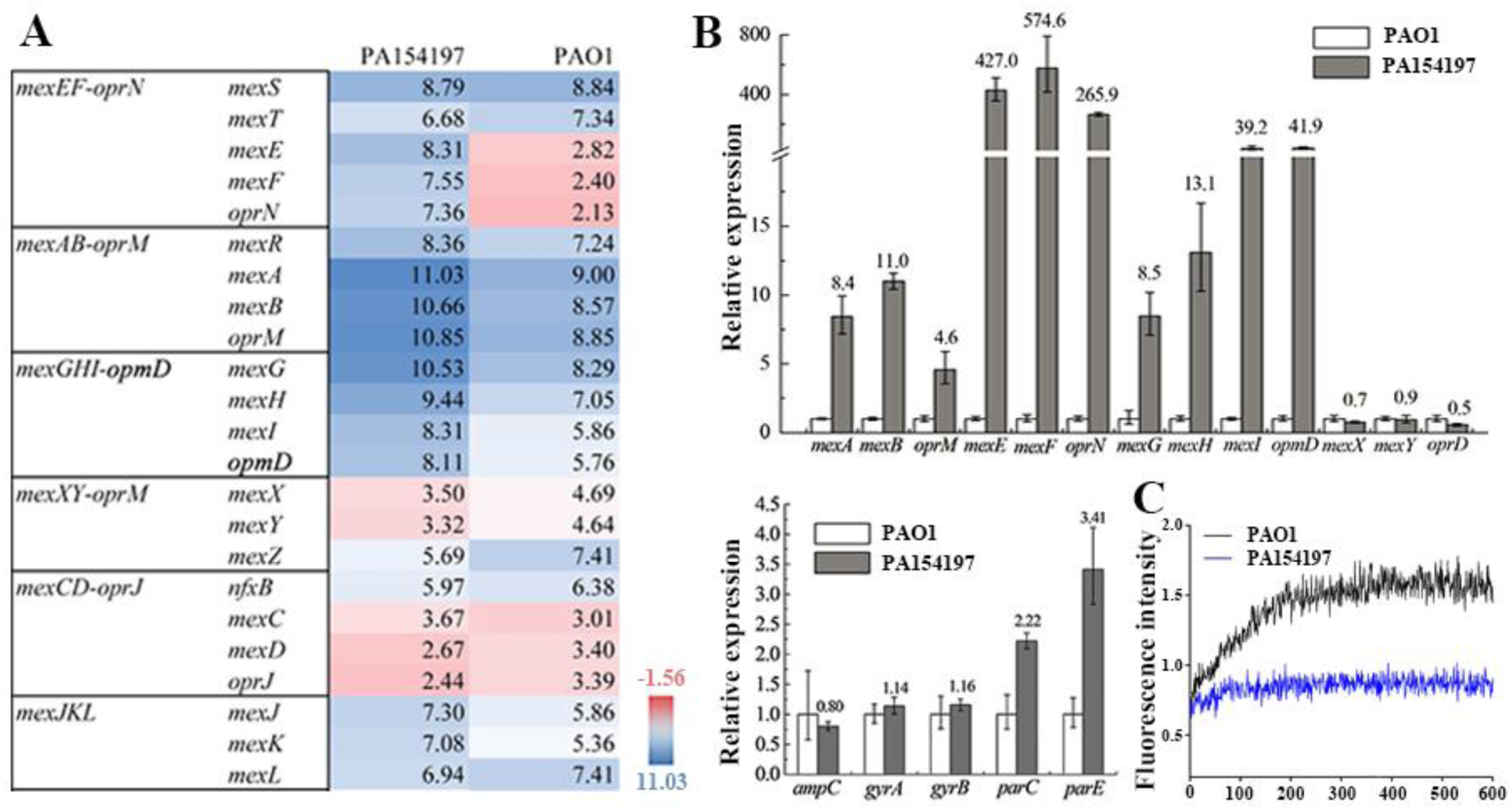
Several multidrug efflux pump genes are over-expressed in *P. aeruginosa* PA154197. **A.** RPKM abundance of the major efflux pump genes in PA154197 and PAO1 calculated from the RNA-Seq data. **B.** RT-qPCR analysis of the expression of selective efflux genes and other genes associated with resistance to specific classes of antibiotics. C. Efflux activities of PA154197 and PAO1 measured by ethidium bromide accumulation. Higher fluorescence indicates lower efflux activity.

### *Virulence of P. aeruginosa* PA154197

Our comparative transcriptome analysis also reveals a significantly higher expression of several virulence genes in PA154197 than in PAO1 (Fig 3 and Table S2). These include the type III (*psc* genes) and the type VI (*tss* and *hcp-1* genes) secretion systems, pyocyanin production (*phz* genes), the PQS quorum sensing system (*pqs* genes), and a series of Fe and heme acquisition genes (*fep* and *has* genes). On the other hand, several genes involved in biofilm formation (such as *psl, alg, lecA*), motility, pili and fimbrial assembly proteins (such as cup genes), and pyoverdine production (*pvd* genes) are expressed at a lower level in PA154197 than in PAO1.

To examine the virulence factors production and the virulence of PA154197, we examined pycyanin (PYO) and pyoverdine production, swimming and swarming motilities, and biofilm formation of PA154197 and compared with that of PAO1. As shown in Fig 5, PA154197 cells display a significantly higher PYO production than PAO1 cells when grown on both LB plate and in LB liquid broth (Fig 5A). In contrast, PA154197 displays reduced pyoverdine production compared to PAO1 (Fig 5B). Consistent with the lower expression of biofilm genes in PA154197 than in PAO1, PA154197 displays a reduced biofilm formation capability as examined by the crystal violet staining (Fig. 5C). In terms of the motility of the bacterium, PA154197 displays a comparable swimming activity with that of PAO1, but a defective swarming motility (Fig. 5D) which is consistent with the lower expression of the pili and fimbria assembly genes in this strain. To evaluate the in vivo virulence of PA154197, we conducted *Caenorhabditis elegans* fast killing and slow killing assays which are established infection models to evaluate the cytotoxicity and pathogenicity of *P. aeruginosa*, respectively(34). We found that the two strains display a comparable cytotoxicity and pathogenicity (virulence) (Fig. 5E) to the *C. elegans* host. Together, these studies indicate that PA154197 over-expresses a subset of virulence factors and does not display a compromised virulence as reported for many other resistant strains.

**Figure 5.**
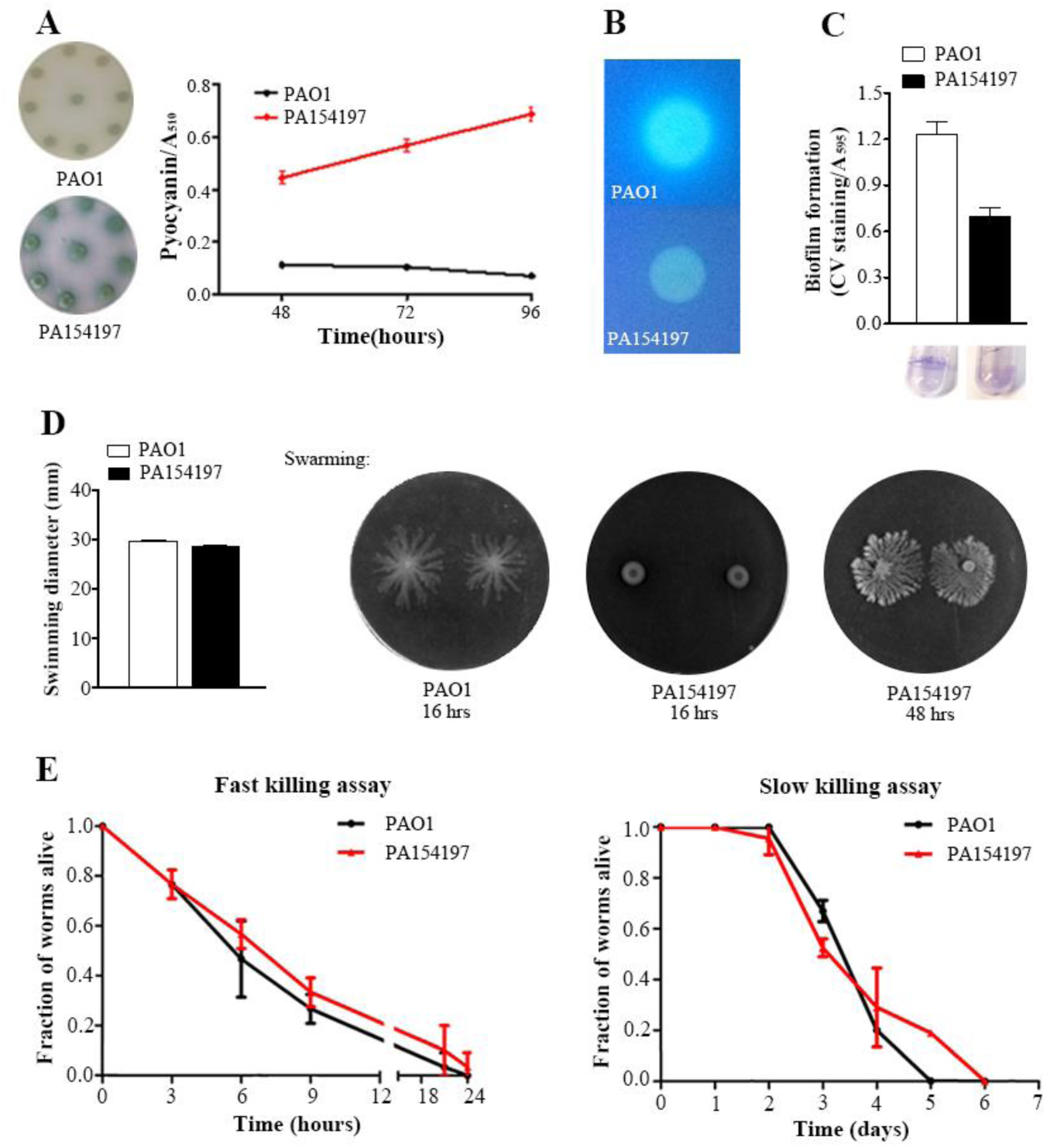
Virulence and production of several virulence factors in PA154197 and PAO1. **A.** Pyocyanin production in PA154197 and PAO1 cells cultured in both LB broth and agar. **B.** Pyoverdine production examined under UV light. **C.** Biofilm formation examined by the crystal violet stain. **D.** Swimming and swarming motilities examined in the corresponding specific agar plates. **E.** Virulence of PA154197 and PAO1 measured in a *C. elegans* infection model. N2 *C. elegans* grown to L4 stage adults were infected with PA154197 or PAO1 for 24hrs. Survival of *C. elegans* were monitored under the light microscope and recorded.

### The Pqs quorum sensing system is activated in PA154197 independent of the primary QS system Las and Rhl

It is known that three hierarchically organized QS systems, Las, Rhl, and Pqs, regulate the production of an arsenal of virulence factors in *P. aeruginosa* with each being primarily associated with the elastase (encoded by *lasB* gene), rhamnolipid (encoded by *rhlA* gene) and pyocyanin (encoded by *phz* genes) production, respectively (35, 36) (Fig 6A). Among them, the Las system proceeds the Rhl and Pqs systems and is the master regulator of the QS circuit. Several previous epidemiological survey and molecular evolutionary studies have identified *lasR* null and mutant variants in clinical isolates and ascribed the attenuated virulence of the isolates to QS deficiency caused by Las inactivation. To investigate the underlying mechanisms of the uncompromised virulence of PA154197, we examined the RNA-Seq transcription levels of the QS systems and their correspondingly regulated genes, especially virulence genes. We found that while expression of the *lasIR* and *rhlIR* in PA154197 is undetectable, expression of the genes encoding the secondary QS system *pqsA-E* is top three significantly expressed (Log2 RPKM abundance in a range of 4.1 – 5.4) gene operon in PA154197 than in PAO1. Consistently, genes primarily controlled by the Las and Rhl systems, such as *toxA, lasB, lecA* are expressed in a lower level in PA154197 than in PAO1, whereas genes controlled by Pqs, such as *phZ* genes, *pqsA-E*, and *phnAB* are expressed in a significantly higher level in PA154197 than in PAO1 (Fig 6A and Table S2). RT-qPCR analysis confirmed this observation (Fig 6B). These data suggest that the secondary QS system Pqs is activated and expressed independent of the primary QS systems Las and Rhl in PA154197, which may account for the hyper-production of a subset of virulence factors and consequently its uncompromised virulence.

**Figure 6.**
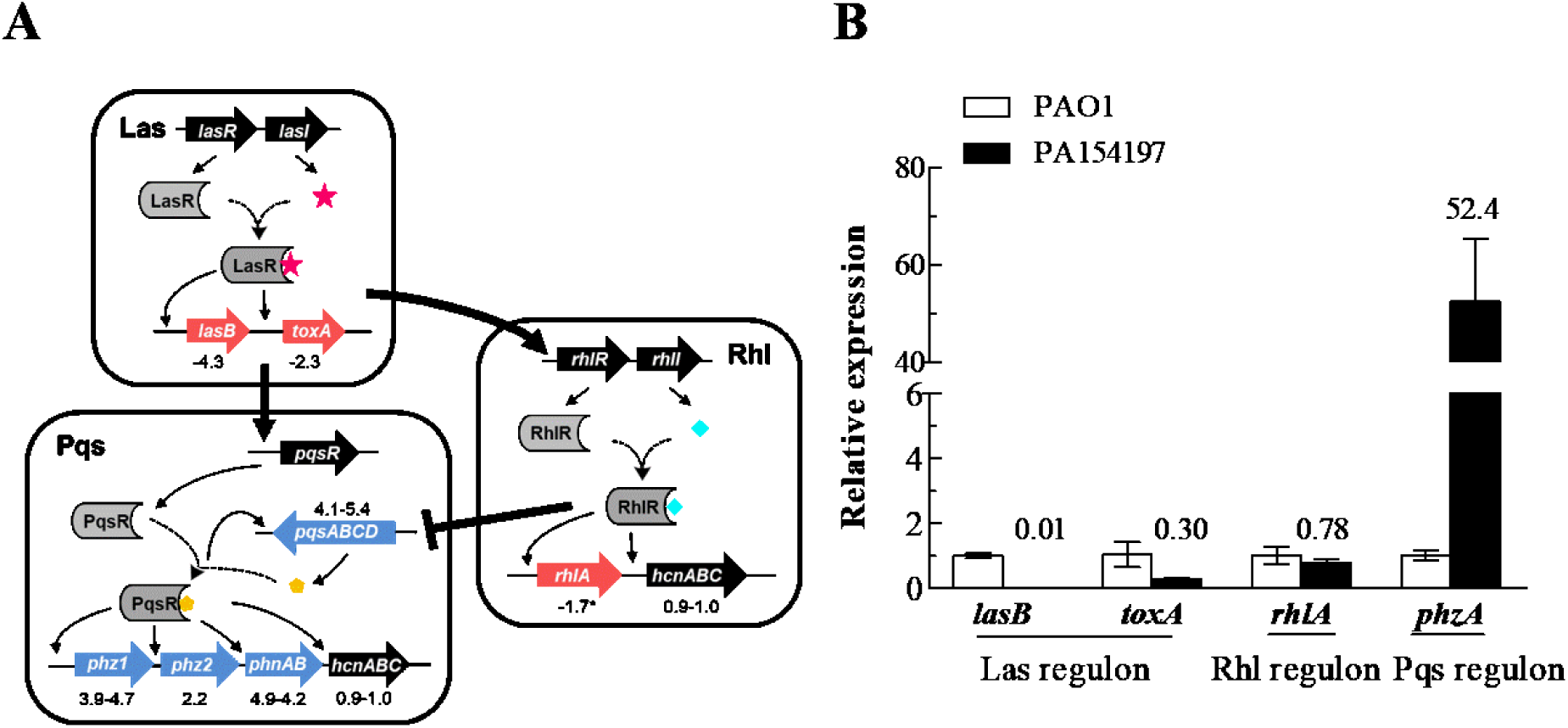
Hyper-activation of the PQS quorum sensing system independent of the master system Las and Rhl in PA154197. **A.** Schematic diagram of the three quorum sensing systems in *P. aeruginosa*, Las, Rhl, and PQS, and their selective regulon. Relative expression of the genes in the Las, Rhl, and PQS regulon is depicted in color scheme with the genes expressed at a lower level in PA154197 than in PAO1 in red color and those expressed at a higher level in PA154197 than in PAO1 in blue color. RPKM abundance of the genes calculated from the RNA-Seq data are indicated. **B.** RT-qPCR of selective genes belonging to the Las, Rhl, and Pqs regulon, respectively.

## Discussion

As a successful and ubiquitous pathogen, *P. aeruginosa* is equipped with extraordinary machineries to adapt to the host environments and antibiotic therapies to survive and disseminate. As a result, the disease development caused by *P. aeruginosa* infections is driven by several dynamic factors, including bacterial pathogenesis, selective forces resulting from the antimicrobial interventions, and the fitness costs of resistance development. It is generally recognized that while acquisition of antibiotic resistance mechanisms confers selective advantages in the presence of antimicrobial therapies, expressing and maintenance of the resistance determinants often incurs metabolic costs to the pathogen and consequently compromises its fitness and pathogenicity potentials (5). How *P. aeruginosa* reconciles this scenario and succeeds in the dynamic cycle of infection and dissemination is not known. In this study, we identify and utilize an MDR *P. aeruginosa* isolate PA154197 which displays a comparable MDR profile to the epidemic high-risk clone ST175 but does not exhibits compromised virulence as a paradigm to examine the underlying pathoadaptive mechanisms. Comparative transcriptome and RT-qPCR analysis reveals an uncoordinated, significant activation of the Pqs quorum sensing system and its regulated virulence genes in PA154197 independent of the primary QS Las and Rhl systems, providing a compensatory mechanism for virulence factor production without the activation of the master and primary QS systems Las and Rhl, as well as the hundreds of genes regulated by them. A schematic diagram to illustrate this pathoadaptive mechanism is shown in Fig 7.

**Figure 7.**
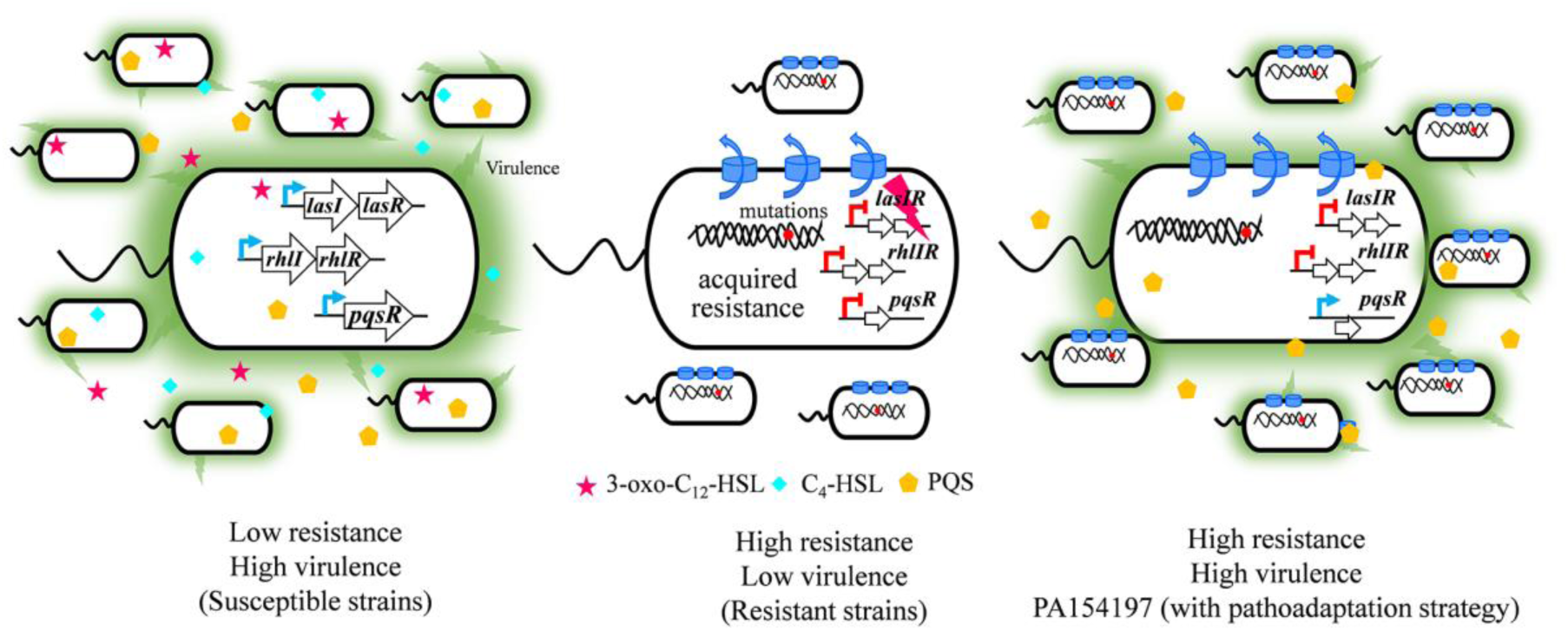
Schematic model to show the pathoadaptation mechanism adopted by PA154197 to modulate its antibiotic resistance and virulence. In the high virulent, susceptible strains, the three quorum sensing (QS) systems Las, Rhl, and Pqs mediate the production of a variety of virulence factors and a high virulence in the cell. Antibiotic resistant strains with acquired resistance determinants often carry mutations in the master QS regulator Las to downregulate the QS system and virulence. Strains with pathoadaptive strategy such as PA154197 uncouple the activation of the secondary QS system Pqs from Las and Rhl to maintain an uncompromised virulence with simultaneous production of its resistance traits.

Several previous studies reported QS deficiency in antibiotic resistant clinical isolates and attributed the attenuated virulence of the strains to this mechanism, as these mutants are proposed to be social cheaters that exploit shared QS products without incurring metabolic costs to themselves (18-21). However, majority of these studies were based on comparative genomic analysis to identify mutations in the QS system genes, and the identified variations were mainly found in the master QS regulator LasR, including both *lasR*-null and various *lasR* point mutations. Consistently, the identified strains often display an overall QS-inactivation phenotype, such as decreased production of all the major QS regulated virulence factors including elastase, rhamnolipids, pyoverdine, pyocyanin. Different from these *lasR* defective isolates, PA154197 produces a significantly higher amount of PYO, a major virulence factor secreted by *P. aeruginosa* (37), and other Pqs regulated gene products than PAO1, while a lower level of Las and Rhl regulated virulence factors and genes than in PAO1. This uncoupled activation of Pqs system ensures the production of a major virulence factor PYO with minimal metabolic burden, thus is beneficial to the fitness and virulence of the pathogen. How *P. aeruginosa* achieves this differential activation is not known currently. We compared the promoter regions of the *lasRI, rhlRI, pqsR*, and *pqsA-E* operons with the corresponding sequences in PAO1, but did not find any nucleotide alternations that may account for the differential activation of *pqs* genes. Comparative genomics analysis revealed a point mutation Q98P in PA154197 LasR which is located at the α5 of the N-3-oxo-dodecanoyl-homoserine lactone binding domain of the protein but the residue is not located in the binding pocket (38). Whether this point mutation leads to differential activation of the Pqs system warrants further investigations. Notably, Las independent activation of a downstream QS system was also reported in a collection of *P. aeruginosa* isolates from CF patients in which half of the *lasR-*null strains were found to retain the RhlR activity (20). Hence, adaptable QS hierarchy which uncouples a downstream QS system from the master regulator LasR may represent an emerging compensatory mechanism that facilitates the fitness, virulence, and persistence of *P. aeruginosa* in the host settings.

With the increasing frequency of detecting the MDR *P. aeruginosa* isolates from various sources, it is recognized that the resistance traits of a strain are often the result of a complex interaction of several cellular processes and no individual mutation or resistance gene is sufficient to confer clinically significant resistance (39, 40). In this study, we identified multiple genetic variations potentially associated with the resistance development in PA154197. These include both machineries which confer resistance to a diverse class of antibiotics and hence causing MDR(4), such as over-expression of multidrug efflux pumps, and genes which mutations result in resistance to specific classes of antibiotics, such as mutations in *ampC* and *gyrA*. Transcriptome and RT-qPCR confirmed the hyper-expression of three multidrug efflux pumps, MexAB, MexEF, and MexGHI in PA154197, but the expression of *ampC* and *gyrA* was found to be in a similar level to that in PAO1. Whether these two genes contribute to the profile and level of resistance exhibited in PA154197 remains further molecular validation. Notably, both the T105A mutation in AmpC and T83I mutation in GyrA are the well characterized genetic variants associated with antibiotic resistance, which have been termed as the PDC-3 type AmpC variant (28) and the quinolone resistance-determining region (QRDR) hotspot mutation, respectively (14). However, these two mutations are also identified in other *P. aeruginosa* isolates we surveyed such as PA150577 which does not display resistance to β-lactams or fluoroquinolones (data not shown). Hence, the observed resistance to antipseudomonal aztreonam (monobactam) and imipenem (carbepenam) in PA154197 may not be due to the T105A mutation in AmpC, and the observed resistance to ciprofloxacin and levofloxacin (fluoroquinolone) may not be due to the T83I mutation in GyrA. In addition, acquiring additional and extended-spectrum β-lactamase on mobile genetic elements represents another common mechanism of β-lactam resistance in *P.aeruginosa* (41), but no plasmid encoding β-lactamases are identified in PA154197. Indeed, it has been proposed that genetic background of the drug resistant strains influences the epistatic interactions of the various resistant determinants and the resistance readout (42). This highlights the intricacy of the mechanisms that underlie resistance development of MDR strains and our ongoing targeted molecular investigations and evolutionary trajectory analysis may provide mechanistic insight into these processes.

## Materials and methods

### Minimum inhibitory concentration measurements

MICs were measured following the standard protocol from ASM with slight modification (43). Single fresh colonies of PA154197 and PAO1 were inoculated in Lysogeny broth (LB) overnight at 37°C with 220-rpm agitation. The resulting cell culture was diluted and distributed to the wells of 96-well plate with a final cell density as 5×10^5^ CFU/ml. Selected antibiotics were added to the wells with concentration ranging from 0.25 to 128 μg/ml. Plates were incubated at 37 °C for 16-20 h. MIC values were determined as the concentration of antibiotics. Antibiotic susceptibility profiles of the strains indicated are displayed in color scheme with low susceptibility (high MIC values) in red color and high susceptibility (low MIC values) in blue color. The color scheme is constructed using background filling application in the Excel of MS Office.

### Genomic DNA extraction

Extraction of the genomic DNA of PA154197 was performed following the description in a previous study (44). Briefly, PA154197 was cultivated in Luria-Bertani (LB) broth overnight with shaking (220 rpm) at 37°C. Bacterial cells were harvested from 1 ml culture via centrifugation at 10,000 rpm for 10 minutes. Genomic DNA was extracted using the QIAamp DNA Mini Kit following the manufacturer’s instructions (Qiagen, Hilden, Germany). The concentration and quality of genomic DNA was determined by NanoDrop and agarose (0.8%) gel electrophoresis.

### Genome sequencing and annotations

Genome sequencing of *P. aeruginosa* PA154197 was conducted on the Illumina NextSeq (300 Cycles) PE150 High Output Flow Cell platform in Georgia Genomics Facility at University of Georgia, USA. SPAdes (45) was used to assemble the reads after removal of adapter, primers, and low quality bases using Trimmomatic (46). The initial assembly of PA154197 genome yielded 21 contigs. The 21 contigs were aligned against the reference genome of PAO1 and ATCC 27853 and gaps were filled using Sanger sequencing to generate the complete genome sequence of PA154197. Gene calling and annotation were carried out using the National Center for Biotechnology Information (NCBI)’s Prokaryotic Genome Annotation Pipeline 2.0 (PGAP) (47).

### Sequence type and phylogenetic analyses

The multiple locus sequence type (MLST) and serotype of PA154197 was predicted based on the PubMLST database (www.pumlst.org) and the *Pseudomonas aeruginosa* serotyper (PAst) tool (26).

To carry out the phylogenetic analysis, we downloaded all available complete genomes of *P*. *aeruginosa* (one representative per distinct phylogenetic group) from NCBI GenBank (78 genomes by Oct 2017) (Table S3, for reviewers’ information only) and selected 72 representative non-complete genomes that are distributed in the 32 phylogenetic groups as defined in the NCBI. Single nucleotide polymorphisms (SNPs) were collected using Parsnp with default parameters (48) and that of PAO1 served as the reference. SNPs were then used to build a maximum likelihood tree in MEGA (49) with the following parameters: Tamura-Nei substitution model, gamma rate distribution among sites, Nearest-Neighbor-Interchange for tree inference options, a bootstrap value 100, and initial tree was generated using the Neighbor Joining method. Variants were called using the Harvest tools (48) and annotated using SnpEff (50).

### Identifications of antibiotic resistance genes (ARG) and mutations therein

Resistance Gene Identifiers (RGI) from the Comprehensive Antibiotic Resistance Database (CARD) (27) were used to identify antibiotic resistance genes in PA154197 using “Perfect and Strict hits only” parameter. BLAST of all genes from the present study against the antibiotic resistance gene database of ResFinder (version 2.1) (51) and BLAST against a database of antibiotic resistance genes curated by ourselves based on previous reports (52) were also conducted.

Variants of all predicted antibiotic resistance genes were collected from the annotation with SnpEff (50) using PAO1 as the reference. SNPs that cause non-synonymous mutations and gaps less than 6 bp were also identified and summarized.

### RNA-seq

RNA extraction, quality control, and RNA-Seq were performed in PA154197 and the reference strain PAO1 with three biological replicates following our previous descriptions (44). Stranded libraries for all RNA samples were constructed using Kapa Biosystems RNA library preparation chemistry in Georgia Genomics Facility at University of Georgia.

Orthologous genes between PAO1 and PA154197 were obtained by comparison using *progressiveMauve* with default settings (32) and were employed in the following RNA-seq analysis. RNA-seq reads were pre-processed for quality control using Trimmomatic (46) and were then mapped to the reference genomes of PAO1 and PA154197 respectively, using Stampy (to speed up the alignment process, alignment was initially conducted using BWA-MEM with default parameters (53) then passed to Stampy with the flags --bamkeepgoodreads -M) (54). SAMtools and BamTools were used for format conversions, statistics, and quality assessment and control (55). The Integrative Genomics Viewer was used to visually inspect mapping quality (56). Fragments counting per genomic features (genes) were performed using featureCounts (*featureCounts -R -M -Q 10 -p -P -s 2 -t gene -g locus_tag --largestOverlap*) (57). Reads that mapped with MAPQ scores below 10 were removed. Enforcing a MAPQ score below 10 also excludes multi-mapped reads albeit the percentage of this category is low (data not shown). DESeq was employed to analyze differentially expressed genes. Selective genes with high expression levels in PA154197 were verified using RT-qPCR. Primers used in the present study are listed in Supplementary Table S4.

### Motility assays

The assays were performed as previously described with slight modifications(58). (1) Swimming motility. The semi-solid motility plates were prepared by mixing the LB broth with 0.25% (wt/vol) agarose. Overnight cultures of PAO1 and PA154197 were sub-cultured (1:200 dilution) into LB broth. When OD_600_ reached 0.1-0.2, 2 μl of cell was spotted at the center of a freshly prepared semi-solid swimming plate. The plates dried at room temperature for 1 h and subsequently were incubated at 37°C for 8 h. Swimming motility was evaluated by measuring the diameter of the covered areas. All assays were conducted in triplicates. (2) Swarming motility. The motility plates were prepared by mixing the M8 broth [21.1 mM Na2HPO4, 11 mM KH2PO4, 42.8 mM NaCl, 9.3 mM NH4Cl, 1 mM MgSO4, 0.2% glucose, 0.2% casamino acids, pH 6.5] with 0.5% (wt/vol) agarose. 2 μl of sub-cultures of PAO1 and PA154197 (OD_600_ of 0.1-0.2) as mentioned above was spotted at the center of a pre-dried swarming plate. The plates were incubated at 37°C, and images of the developed swarms were recorded at 16-48 h. All assays were conducted in triplicates.

### Pyocyanin production assay

Pyocyanin production was examined according to a protocol described previously with modification (44). 1 ml of the bacterial culture was subject to centrifugation at 13,000 rpm for 5 min. The supernatant was collected and extracted with 600 μl chloroform following vortex for 10 s twice. After centrifugation at 13,000 rpm for 5 min, the chloroform phase was transferred into a clean tube, and subsequently mixed with 0.5 ml 0.2 M HCl followed by gentle shaking to transfer the pyocyanin to the aqueous phase. The concentration of PYO was determined by measuring the absorbance of the aqueous phase at 510 nm.

### Biofilm assay

Crystal violet staining method was used to determine biofilm formation of PAO1 and PA154197 with slight modifications (59). Overnight cultures of *P. aeruginosa* cells were inoculated into 0.5 ml of LB broth in 5 mL round bottom polypropylene Falcon tubes (final cell density as 0.1) and incubated statically at 37 °C for 24 or 48 h to allow biofilm formation. After removing the medium, the biofilms formed at the bottom of the tubes were gently washed with phosphate buffered saline (PBS) three times to remove residual planktonic cells. The tubes were then air-dried and stained with 0.1% crystal violet for 15 min at room temperature. After washing the tubes three times with sterile distilled water to remove excess dye molecules, biofilms were dissolved in 150 μl of 30% acetic acid and the optical density (OD) was measured at 595 nm using a 96-well plate spectrometer reader. All experiments were performed in triplicates.

### Nematode killing assay

To examine the virulence and pathogenesis of PA154197 compared with PAO1, fast and slow killing kinetics of *Caenorhabditis elegans* (*C. elegans*) were performed with slight modification (60). (1) Fast killing assay. Preparation of the killing plates can be divided into two steps. i) 100 μL of overnight culture of PA154197 or PAO1 were spread onto 3.5 cm LB agar plates and incubated at 25°C for 2 days to produce the toxins. ii) Lawn of PA154197 or PAO1 was scraped from LB agar plates containing the toxins completely using a sterile L spreader. *C. elegans* strain N2 were synchronized by isolating eggs from gravid adults and plating eggs on to lawns of *E. coli* OP50 on NGM [0.25% peptone, 0.3% NaCl, 2% agar, 5 μg/ml cholesterol, 1 mM MgSO4, and 25 mM KH2PO4 (pH 6)] agar plates followed by incubation at 25°C for 48-52 h to reach the L4 stage. Each fast killing plate was seeded with 20-30 worms. Plates were incubated at 25°C and scored every 4-6 h. A worm was considered to be dead when it no longer responded to touch. (2) Slow killing assay. The procedure was similar as fast killing assay except the preparation of killing plates. For slow killing plates, lawn of PA154197 or PAO1 was not moved away. All experiments were performed in triplicates.

### Pyoverdine (PVD) production assay

The siderophore pyoverdine is observed under UV light as a fluorescent zone around the colony (61). Fresh single colonies of PA154197 and PAO1 were inoculated in LB medium overnight at 37 °C with 220-rpm agitation. Following diluting overnight culture to 1×10^9^ CFU/ml with fresh LB medium, 10 μl cell suspension was then spotted on a LB agar plate. The plate was incubated at 37 °C for 16-20 h. Fluorescent PVD was visualized and imaged under UV light.

### Efflux activity assay

Efflux activity of PA154197 and PAO1 cells is examined by ethidium bromide accumulation as described previously (62). Overnight cultures of PA154197 and PAO1 cells were diluted 1:100 in fresh LB broth and grown at 37°C until the OD_600nm_ reached 0.5-0.6. The cells were collected by centrifugation at 4,000g for 10 min at 4°C, washed three times with efflux buffer (50 mM potassium phosphate buffer, pH 7.0, containing 5 mM MgSO4), and then resuspended in the same buffer to an OD_600 nm_ of 0.5. Following incubation at room temperature (25°C) for 3 min in the presence of 25 mM glucose to energize the cells, ethidium bromide (Sigma-Aldrich) was added to a final concentration of 2 μM. The ethidium bromide fluorescence was measured continuously using the Varioskan Flash Multimode Microplate Reader (Thermal Scientific) at Ex/Em wavelengths of 500 nm/580 nm. Higher fluorescence signal indicates lower efflux activity. Assays were performed in triplicates and mean values were used to plot the efflux curve.

## Acknowledgements

We thank Dr. Karen Yuen (School of Biological Sciences, HKU) for her help to establish the *C. elegans* killing assay. This work is supported by the Hong Kong University Grants Council General Research Fund (17142316, to AY), Seed Funding for Strategic Interdisciplinary Research Scheme (HKU 2017, to AY), and Shenzhen City Knowledge Innovation Plan (JCYJ20160530174441706 to AY).

## Author Contributions

HC, AY designed the studies. HC, TX, YL, ZX, YKL, SB, VBB, conducted the experiments. HL provided sequencing service. PW provided clinical strains. HC, TX, YL, ZX, AY wrote the manuscript.

The authors do not have conflict of interests to declare.

